# Grading HER2 at the nanoscale in clinical tissue

**DOI:** 10.64898/2026.01.14.695095

**Authors:** Lauren K. Toms, Hervé Barjat, Isabel Peset, Tiffany J. Allen, Kevin Randall, Denis G. Alferez, Robert B. Clarke, Emily P. Offer

**Affiliations:** Medicines Discovery Catapult, Block 35, Mereside, Alderley Park, Macclesfield, Cheshire, SK10 4ZF; Spanish National Cancer Research Center, Madrid, Spain; Breast Biology Group, Division of Cancer Sciences, School of Medical Sciences, Faculty of Biology, Medicine and Health, University of Manchester

**Keywords:** single-molecule localisation microscopy, HER2, breast cancer, clustering, super-resolution microscopy, STORM, Earth Mover’s Distance

## Abstract

To guide diagnosis and treatment, breast cancer biopsies are assessed for HER2 status and assigned one of four grades (0-3+). While current practices are sufficient for detection of HER2 overexpression (3+), there is a need for more sensitive methods capable of characterising lower HER2 expression in patients who may still benefit from HER2-targeted therapies. Super-resolution fluorescence microscopy techniques, such as single molecule localisation microscopy (SMLM), have reshaped the study of nanoscale molecular architecture by visualising single target molecules in a range of sample types. Here, we have developed a quantitative SMLM workflow to visualise HER2 nanoclustering in patient-derived xenografts (PDX) and clinical breast tumour tissue from eight patients spanning all disease grades. Analysis of HER2 cluster architecture revealed grade-dependent changes in size of cluster and HER2 abundance. We then applied a blinded data-driven approach to regroup samples based on this nanoscale HER2 clustering. This led to the reclassification of three samples into new groups, due to similarities in nanoscale signature. Together, these findings demonstrate that quantitative fluorescence nanoscopy can be used to identify clinical HER2 phenotypes across a range of expression levels due to its exquisite sensitivity, and this could be leveraged to stratify patients for targeted therapy.

## From conventional scoring to nanoscale insight: super-resolution microscopy as a tool for HER2-targeted therapy stratification

Breast cancer is a significant global burden, affecting 2.3 million women and causing 670,000 deaths in 2022 alone ^1^. The receptor tyrosine kinase, Human Epidermal Growth Factor Receptor 2 (HER2), is overexpressed in ∼15-20% of breast cancers and is associated with a poor prognosis and aggressive phenotype if left untreated ^2^. However, anti-HER2 therapies, such as the monoclonal antibody trastuzumab, have revolutionised the treatment of HER2-positive breast cancer and greatly improved outcomes for patients ^3^. Administration of anti-HER2 therapy is dependent on the status of HER2 expression, which is initially assessed by immunohistochemistry (IHC) of a biopsy sample. Based on the degree of HER2 membrane staining, the tumour is assigned one of four grades (0, 1+, 2+ or 3+) by a clinician ^4–7^. A grade of 0 or 1+ is classed as HER2 negative, while 3+ is HER2 positive and will proceed to anti-HER2 therapy ^5^. HER2 2+, also known as equivocal, tumour samples require further testing by in situ hybridisation (ISH) to quantify HER2 gene amplification and are then reclassified as HER2 positive or negative ^5^. A small percentage of patients are ISH equivocal and require additional testing to identify the treatment route ^5^. Recent clinical trials have questioned whether this classification sufficiently captures HER2 status in the context of new therapeutics. Indeed, the DESTINY-Breast04, DESTINY-Breast06 and DAISY trials have found that patients with HER2 ‘low’ (IHC 1+ and 2+ without gene amplification) and ‘ultralow’ (a subset of IHC 0) tumours can benefit from a HER2-targeted antibody-drug conjugate (ADC) called trastuzumab deruxtecan (T-DXd) ^8–10^. Although this expands treatment options, the assignment of HER2 low and ultralow status is challenging, treatment route unclear, and research is ongoing. Current guidelines, without adopting the HER2 low and ultralow terminology, now recommend best practice efforts to distinguish between IHC 0 and 1+ HER2 grades, however, acknowledge the risk of suboptimal sensitivity of the traditional IHC diagnostic assay to detect low levels of HER2 ^5^. Additionally, a low concordance rate between pathologists in differentiating between IHC 0 and 1+ tumours has been reported ^11,12^. As novel therapeutics emerge and technology advances, the points above highlight a clear need for more sensitive diagnostic assays for patient stratification.

Super-resolution microscopy is an umbrella term for techniques that improve the resolution of fluorescence microscopy beyond a theoretical barrier known as the diffraction limit ^13^. Despite being developed approximately 20 years ago, its application in drug discovery and diagnostics has only recently been experimentally considered ^14^. Single-molecule localisation microscopy (SMLM) approaches such as direct stochastic optical reconstruction microscopy (dSTORM) improve resolution by localising few, spatially separated fluorophores per time frame and integrating localisations acquired from many thousands of frames into a final image with spatial and temporal information about the detected localisations. This increases resolution by an order-of-magnitude to < 30 nm and enables the nanoscopic investigation of many sample types with visible light, which has been particularly impactful in the study of receptor complexes ^15,16^. The enhanced resolution these techniques enable may be key to understanding disease phenotypes with their utility already evidenced in clinical tissues including in breast cancer samples ^17–19^, renal tissue ^20^ and colorectal tumours ^21,22^.

Receptor clustering is a common physiological and pathological phenotype, which is suggested to underpin cellular function by mediating a rapid response to stimuli, stabilising signalling centres, and increasing receptor specificity ^23^. In the case of HER2, research has found that high expression leads to the formation of elongated HER2 clusters along the cell surface, which induce membrane deformation and may contribute to an invasive phenotype ^24^. Moreover, the organisation of HER2 is dynamic in response to the therapeutic antibodies trastuzumab and pertuzumab ^25,26^, which deplete HER2 from the plasma membrane and induce a clustering phenotype ^27,28^. This is of importance as the density and clustering of HER2 is suggested to drive trastuzumab sensitivity in both cell lines and patient biopsies ^18,25^. Altogether, these works demonstrate the relationship between HER2 clustering and pathophysiological response and highlight the potential of SMLM to guide clinical decision-making through quantification of HER2 nanosignatures.

In this study, we have used dSTORM imaging and cluster analysis to investigate the organisation of HER2 at the nanoscale in breast cancer tissue from patients spanning all clinical grades. We find that HER2 nanosignatures are inconsistent within grades and apply a blinded data-driven reclassification to group samples based on this fundamental molecular feature. We suggest this could inform new stratification methods that leverage the sensitivity of localisation microscopy to further personalise anti-HER2 therapy, which may be of particular use for low expressing patients.

## Results and Discussion

### Quantification of HER2 nanoclustering in PDX samples

Prior to undertaking work in clinical samples, we sought to develop a method to quantify HER2 nanoclustering using dSTORM in formalin-fixed paraffin-embedded (FFPE) PDX samples. Five PDX samples (P1-5) were assigned an IHC HER2 grade by a research pathologist using the guidelines for diagnosis ^4–7^. Representative images of the PDX samples labelled for HER2 and imaged by IHC and immunofluorescence (IF) widefield microscopy are shown in Figure 1a and 1b, respectively, and staining controls depicted in Supplementary Figure 1. Both techniques qualitatively show concordance for all samples. P1 (IHC 0) has little to no HER2 staining. P2 (IHC 1+) has weak membrane staining. P3 (IHC 2+) shows moderate membrane staining and P4 and P5 (IHC 3+) show strong membrane staining.

**Figure 1:**
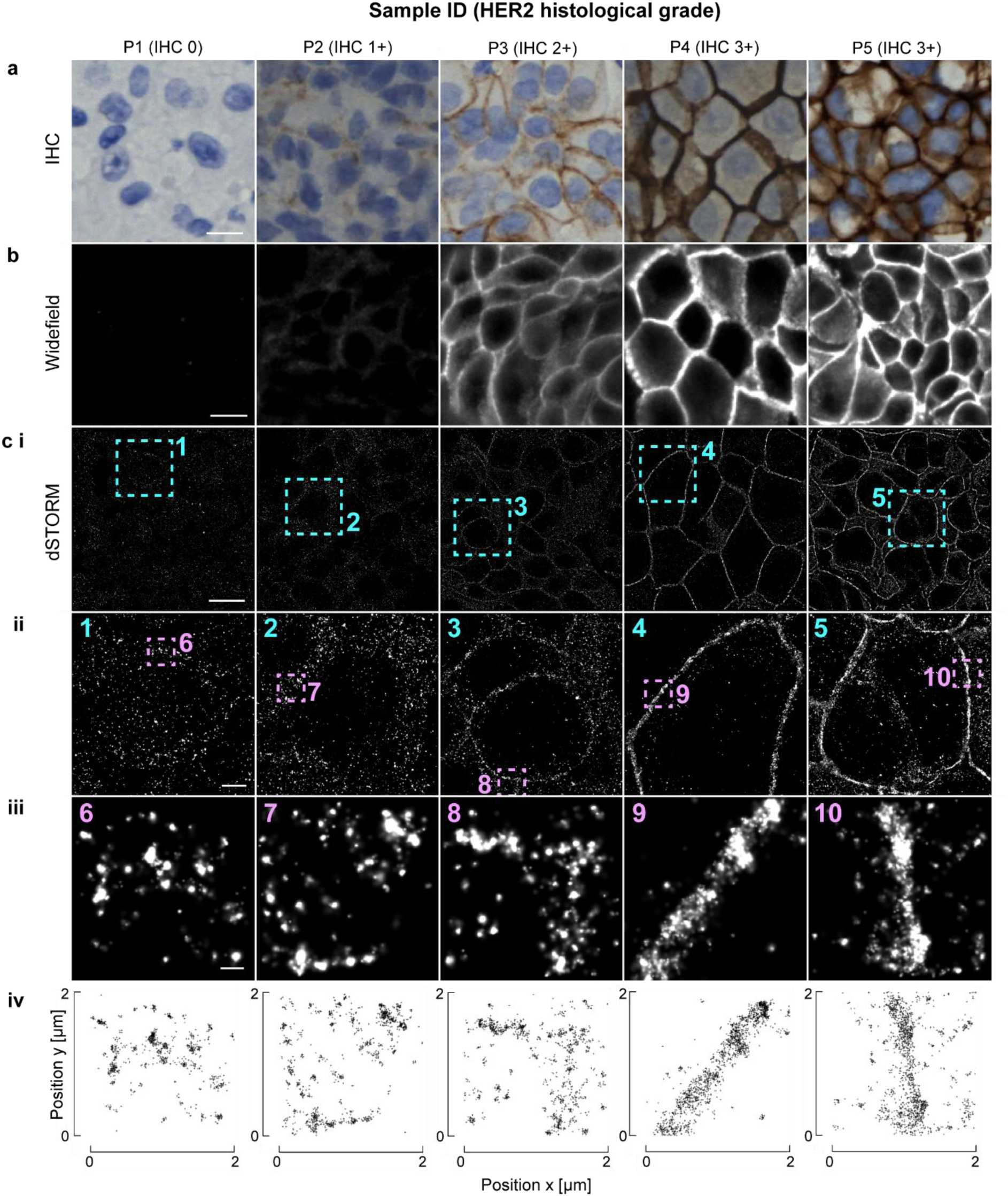
Visualisation of HER2 protein in five patient-derived xenografts (P1-5) using IHC (a), widefield fluorescence microscopy (b), and dSTORM (c). dSTORM images in ci to ciii are shown with a 2D gaussian-render, with progressive zoomed regions shown in cii and ciii. The coordinates of the detected localisations in ciii are shown in civ. HER2 grading was assigned using immunohistochemistry (IHC 0-3+). Scale bars: 10 µm (a, b and ci), 2 µm (cii) and 250 nm (ciii).

To understand HER2 nano-organisation, each PDX sample was visualised using dSTORM and images reconstructed from 5,000 frames (Figure 1c). In agreement with IHC and IF, dSTORM identified progressive membranous abundance of HER2 with IHC grade (Figure 1ci-iii). Furthermore, HER2 localisations in the 3+ samples visualised by dSTORM appear as discontinuous foci, consistent with the notion that HER2 adopts a clustered organisation ^18,19^ (Figure 1ciii). These apparent clusters were also observed in the IHC and IF negative P1, demonstrating how the increased sensitivity of dSTORM enables the detection of HER2 ultralow status (Figure 1ciii).

To quantify this HER2 clustering phenotype in an automated fashion, we used the clustering algorithm Topological Mode Analysis Tool (ToMATo, see methods) and assigned the dSTORM localisations into clusters (Figure 2ai,aii) ^29^. We found that while the number of clusters, their area, and density remained constant between grades, the number of localisations per cluster was greater in the 3+ samples when compared to 2+, 1+ and 0 (p = 0.002, p < 0.001 and p < 0.001, respectively) (Figure 2bi-biii, Supplementary Figure 2b). Moreover, quantification of the total localisations per tissue area revealed a progressive increase from grade 1+ to 3+, while no detectable difference was observed between grades 0 and 1+ (Supplementary Figure 2a). Since localisations are directly proportional to the abundance of HER2 molecules, this is consistent with overexpression and the increase in fluorescence intensity observed in the diffraction limited images (Figure 1b, Supplementary Figure 2a). Altogether, these data demonstrate that HER2 is organised into nanoclusters in PDX samples, and these clusters have distinct features that change with clinical grade. Nevertheless, PDX samples should be used with caution as they can misrepresent clinical tissue due to differences in host environment, tumour heterogeneity, and/or selection pressure during PDX propagation ^30^. Therefore, to draw conclusions about the potential clinical utility of SMLM-based HER2 analysis, we must assess the receptor nano-organisation in primary samples acquired in the clinic.

**Figure 2:**
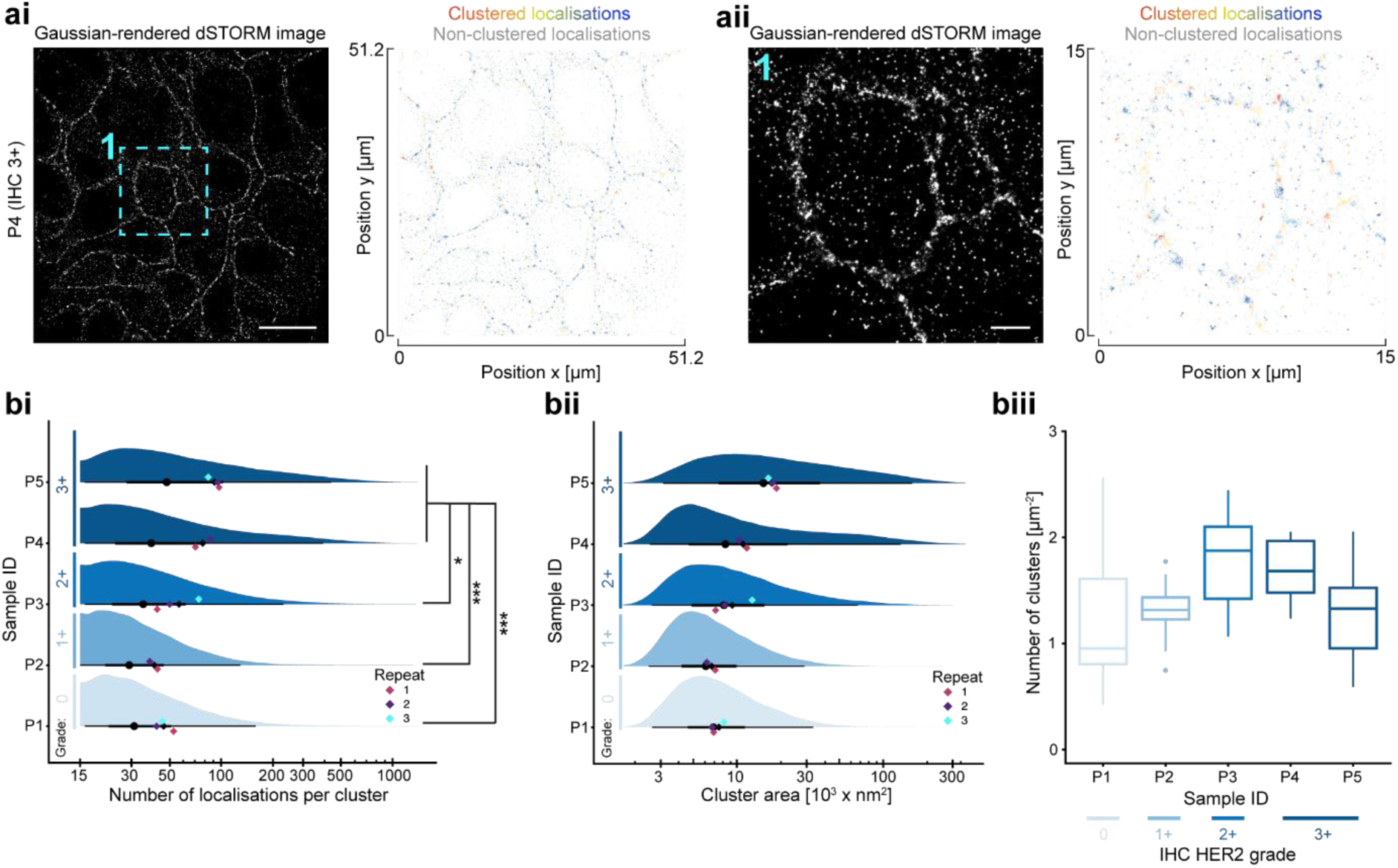
HER2 cluster analysis in five PDX samples (P1-5). (a) Example localisation assignment into clusters using clustering algorithm ToMATo^29^. The gaussian-rendered dSTORM image is shown next to the assignment image with clustered localisations shown in colour and unclustered localisations in grey. A zoomed region from ai is displayed in aii. Scale bars: 10 µm (ai) and 2 µm (aii). The number of localisations per cluster (bi), cluster area (bii) and number of clusters per tissue area (biii) were quantified. The sample mean (black diamond) and median (black circle) are shown with lines representing the interquartile range (thick line) and the range of 95% of the data (thin line) (bi and bii). HER2 grading was assigned using immunohistochemistry (IHC 0-3+). N = 39,731-82,374 clusters from 2-3 technical repeats. Significant differences were determined using a Linear Mixed-Effects model with Tukey HSD test. p ≤ 0.05 (*), p ≤ 0.001 (***).

### dSTORM of HER2 in clinical tissue reveals grade-independent clustering phenotypes

To investigate the conservation of HER2 nanoclustering phenotypes in clinical tissue, we reconstructed dSTORM images acquired with 5,000 or 10,000 frames from eight patient tumour samples (S1-8) identified by Haematoxylin & Eosin (H&E) staining (Figure 3a, Supplementary Figure 3). Samples were graded 0-3+ by a research pathologist upon receipt from the biobank, with two samples from each IHC HER2 grade available. In agreement with the PDX samples, widefield microscopy showed a progressive increase in anti-HER2 fluorescence intensity in a grade-dependent manner (Supplementary Figure 4). We found clustering properties to be comparable when analysing images reconstructed from 5,000 or 10,000 frames therefore the following analyses are based on images reconstructed from 5,000 frames only (Figure 3, Supplementary Figure 5). Importantly, in this case, acquiring only 5,000 frames increased data acquisition throughput two-fold without losing information, but further validation is required with a larger sample cohort.

**Figure 3:**
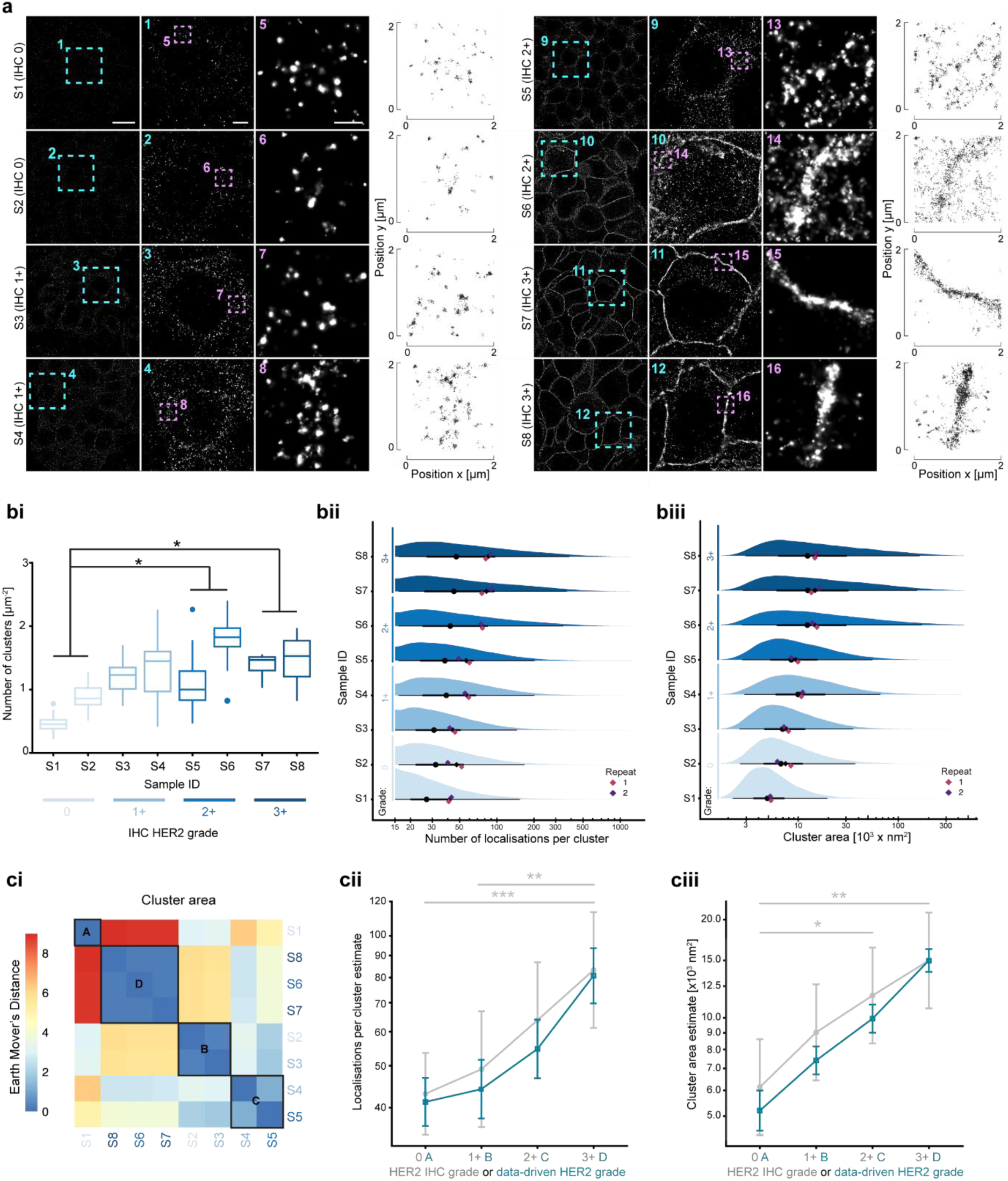
HER2 cluster analysis in eight patient breast tumour samples (S1-8). (a) Example gaussian-rendered dSTORM images of HER2 for each sample. Zoomed regions are shown by the boxes and respective numbering. Individual localisations are shown for the second zoomed image. The immunohistochemistry (IHC) grade is stated next to the sample ID. Scale bars (from left to right): 10 µm, 2 µm and 500 nm. The number of HER2 clusters per tissue area (bi), cluster area (bii) and number of localisations per cluster (biii) were quantified. The sample mean (black diamond) and median (black circle) are shown with lines representing the interquartile range (thick line) and the range of 95% of the data (thin line) (bii and biii). N = 22,817-119,756 clusters from 2 technical repeats. The Earth-Mover’s Distance heatmap (ci) shows the data-driven grouping of the samples into A, B, C and D. Comparison of the mean ± 95% confidence interval for the IHC grading and data-driven grading is shown in (cii) for number of localisations per cluster and (ciii) for cluster area. Significant differences were determined using a Linear Mixed-Effects model with Tukey HSD test. p ≤ 0.05 (*), p ≤ 0.01 (**), p ≤ 0.001 (***). Significant differences between HER2 IHC grades only are shown in cii and ciii.

When considering the mean number of localisations per tissue area (clustered and non-clustered), quantification showed an increase from 43.6 µm^-2^ (95% confidence interval: 35.9, 51.3) in IHC grade 0 to 90.0 µm^-2^ (82.4, 97.5, p < 0.001) in 1+, which continued to 141.7 µm^-2^ (134.2, 149.2) in 2+ (p < 0.001). No difference was detected between grades 2+ and 3+ (156.9 (149.3, 164.4), p = 0.07) (Supplementary Figure 7). These localisations were assigned into clusters using ToMATo as above, which revealed several grade-dependent phenotypes. First, the mean number of clusters was found to only differ when comparing grade 0 with grades 2+ and 3+ (p = 0.04 in both cases, Figure 3bi). Similarly, the cluster area was only found to increase between grade 0 and grades 2+ and 3+ (p = 0.04 and 0.001, respectively, Figure 3biii,ciii). When comparing individual clusters, we found the number of localisations per cluster, in agreement with the PDX samples, was greater in IHC 3+ when compared to grades 0 and 1+, with no other differences observed (p < 0.001 and p = 0.004, respectively, Figure 3bii). Altogether, this analysis of grade-dependent changes in HER2 nanoclustering demonstrates that cluster number and area increase when comparing IHC grade 0 with grades 2+ and 3+, and clusters in IHC 3+ samples contain more HER2 molecules than grades 0 and 1+.

Our data reveal that grade-dependent changes in nanoclustering phenotype are only evident at the extremes, such as when comparing IHC 0 with grades 2+ and 3+. This results from variation within IHC-assigned grade, suggesting this designation might not capture the complexity of HER2 nano-organisation (Figure 3bi-biii). To explore this, we used a blinded data-driven analysis using the Earth Mover’s Distance (EMD) to group samples into four classes based on the distribution of HER2 cluster area independent of IHC grade (Figure 3ci). This new data-driven grouping, summarised in Table 1, isolated S1 (IHC 0) into group A and assigned samples S2 (IHC 0) and S3 (IHC 1+) into group B. Samples S4 (IHC 1+) and S5 (IHC 2+) were assigned to group C, and S6 (IHC 2+), S7 and S8 (IHC 3+) to group D. Following reclassification, a progressive increase in cluster area was observed between all groups, with localisation number following a similar trend except for comparison between A and B (p < 0.001 in all cases, Figure 3cii, ciii). Given that reclassification occurred across IHC grades, this approach might have utility in distinguishing between HER2 ‘low’, HER2 ‘ultralow’ and HER2 ‘null’ as well as classifying HER2 equivocal samples. Future work utilising a larger sample cohort is needed to determine how the HER2 nanoclustering continuum changes with increasing disease severity, enabling groups to be fine-tuned and contextualised with treatment route and clinical outcome.

**Table 1:** HER2 status for each sample assigned by IHC and data-driven analysis of HER2 clustering.

| Sample ID | IHC HER2 grade | Data-driven group |
| --- | --- | --- |
| S1 | 0 | A |
| S2 | 0 | B |
| S3 | 1+ | B |
| S4 | 1+ | C |
| S5 | 2+ | C |
| S6 | 2+ | D |
| S7 | 3+ | D |
| S8 | 3+ | D |

### Summary and future perspectives

In this study we have developed a quantitative SMLM approach to study HER2 nanoclustering in clinical breast cancer tissue spanning all IHC grades. This showed that while a general increase in receptor nanoclustering could be detected between low and high grades, variation was evident within individual IHC grades that obscured the expected progressive HER2 trend. Because of this, we applied a data-driven reclassification approach to group samples based on similarities in their HER2 nanosignature, which yielded four distinct classes of patient sample. While investigations in a larger cohort are required to reaffirm these findings, we suggest the clear grouping observed in this study means the approach could have clinical utility.

We propose this dSTORM-based quantification of HER2 nanoclustering could be applied for personalised intervention. Clinical evidence from studies of anti-HER2 therapy support the expansion of treatment groups. For example, trastuzumab deruxtecan (T-DXd) is suggested to benefit patients with HER2 low and ultralow tumours ^8–10^. As an antibody-drug conjugate, T-DXd is internalised into cells via HER2, deruxtecan is then cleaved to elicit DNA damage and cell death. As a result, deruxtecan can freely diffuse into neighbouring cells and therefore also trigger cell death in cells that are HER2-negative, which may indicate why it shows efficacy in ultralow HER2 status patients ^31,32^. We suggest the sensitivity of dSTORM imaging allows this heterogenous nano-organisation in low expressors to be reliably quantified and ultimately used for stratification. Additionally, trastuzumab resistance is common due to protein truncation and epitope masking ^33^. The presented SMLM workflow could be used to predict patient response by identifying if the clinical antibody binds in the context of HER2 nanoclustering which underlies treatment sensitivity ^18,25^. Importantly, this information can reduce unnecessary health burdens on patients, for example trastuzumab induced cardiotoxicity in patients that will not benefit from trastuzumab ^34^.

The presented approach is not limited to analysis of HER2 nor to breast cancer tissue analysis because protein nanoclusters are widespread in signalling and disease. Since pathogenesis in breast cancer is driven by a wide range of membrane-bound receptors, determining HER2 nanocluster context, for example receptor heterodimer expression with multiplexed imaging is an attractive prospect. Alternative SMLM approaches, such as DNA point accumulation for imaging in nanoscale topography (PAINT)^35^, are capable of this multiplexing, and when combined with similar blinded quantification, could provide a detailed molecular profile of each patient based not only on protein abundance, but proximity and likely interactions. Furthermore, by simply swapping detection reagents the presented approach could provide insight into the nanoclustering of epidermal growth factor receptor (EGFR) overexpressing cancers (e.g., glioblastoma, colorectal and oesophageal cancers), Ras-mutant cancer (e.g., pancreatic, colorectal, lung and melanoma) and even immune and inflammatory diseases ^20,36,37^. The power of such investigation lies not only in better stratification of existing disease states, but in the identification of novel early pathogenic biomarkers ^38^.

With super-resolution microscopy techniques becoming more accessible in terms of equipment size, cost, and operational expertise, their application in a clinical setting is ever closer to realisation. Our data demonstrates the potential of dSTORM for stratification of HER2 nano-phenotypes across a range of expression levels in patient tissue. Broadening this work to a large sample cohort with a focus on clinical outcome will determine how such analysis can further personalise intervention by improving diagnosis.

## Methods

### Ethics

All clinical samples were approved by our internal ethics committee and Manchester Cancer Research Centre Biobank Research Tissue Bank Ethics (ref: 22/NW/0237).

### Samples

All samples used in this study were formalin-fixed paraffin-embedded (FFPE) tissue.

Patient derived xenografts (PDX), provided by Manchester Cancer Research Centre, were used for method development and proof of concept preliminary work ^39^. The PDX samples had varying grades of HER2 which were immunohistochemically assigned by a research pathologist, Table 2.

**Table 2:**
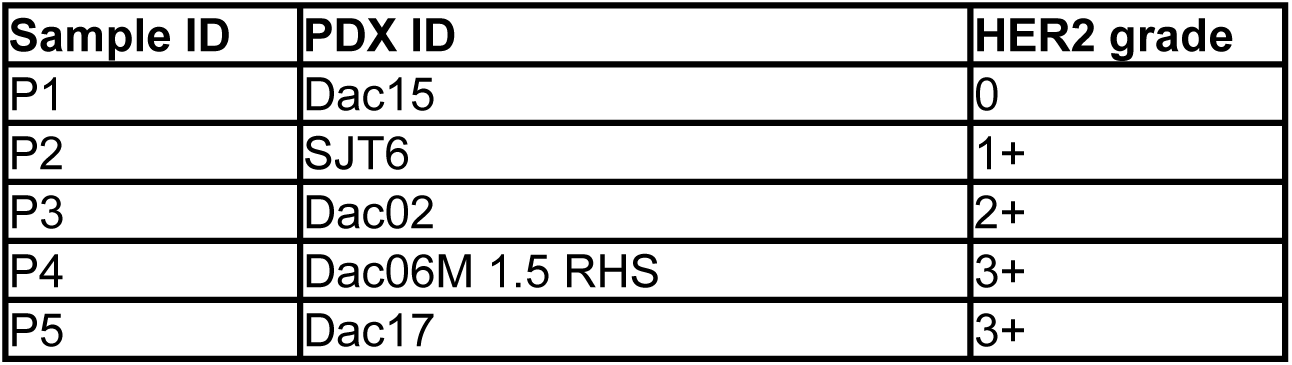
Patient derived xenograft (PDX) samples used in this study. Sample ID (referred as in text), PDX sample ID and immunohistochemical HER2 grade.

| Sample ID | PDX ID | HER2 grade |
| --- | --- | --- |
| P1 | Dac15 | 0 |
| P2 | SJT6 | 1+ |
| P3 | Dac02 | 2+ |
| P4 | Dac06M 1.5 RHS | 3+ |
| P5 | Dac17 | 3+ |

Once the workflow was fully optimised, it was applied to clinical samples of which we had eight samples, Table 3. All clinical samples were provided by Manchester Cancer Research Centre and HER2 grading carried out by a research pathologist.

**Table 3:** Clinical samples used in this study. Sample ID (as referred to in text), Biobank sample ID and IHC HER2 grade.

| Sample ID | Biobank sample ID | HER2 grade |
| --- | --- | --- |
| S1 | BB6RC80 | 0 |
| S2 | BB6RC20 | 0 |
| S3 | BB6RC140 | 1+ |
| S4 | BB6RC130 | 1+ |
| S5 | BB6RC119 | 2+ |
| S6 | BB6RC107 | 2+ |
| S7 | BB6RC87 | 3+ |
| S8 | BB6RC39 | 3+ |

The FFPE tissue blocks were sectioned at 4 µm thickness onto 22 x 22 mm #1.5 high precision coverslips (for immunofluorescence (IF)) or microscope slides (for haematoxylin and eosin (H&E) stain or immunohistochemistry (IHC)). The high precision coverslips were coated with poly-l-lysine (P8920, Sigma) (0.01% v/v in ultrapure water for 10 minutes, then dried at 70 °C for 30 minutes).

Prior to staining and labelling techniques described below, FFPE tissue sections were dewaxed by immersing in xylene (XYL050, Genta Medical) for four minutes, twice. Sections were then rehydrated through graded concentrations of ethanol (100%, 100%, 95%) and finally washed with pure water.

### Haematoxylin and Eosin stain (H&E)

All samples were stained with H&E to identify the tumour cells within the tissue section. A standard protocol was used as follows; slides were immersed in Haematoxylin (10034813, Fisher Scientific) for 2 minutes, followed by 1 minute water, 30 seconds Clarifier (10603639, Fisher Scientific), 1 minute water, 10-15 dips in bluing solution (10862090, Fisher Scientific), 1 minute water, 10 dips in 95% ethanol, 20 seconds Eosin Y (10046868, Fisher Scientific), 10 dips in 95% ethanol, 10 dips in 100% ethanol (twice) and allowed to dry for 10 minutes before mounting onto coverslips with ClearVue® Mountant XYL mounting medium solution (12766788, Fisher Scientific).

### Antigen retrieval

Before carrying out immunolabelling for HER2, the tissue sections underwent heat induced antigen retrieval. The sections were immersed in 1 mM EDTA buffer for 20 minutes at 95 °C using a Milestone KOS microwave.

### Immunohistochemistry (IHC)

IHC was carried out on the PDX samples only. Staining was carried out using a Leica BOND RX automated staining workflow with Leica BOND reagents, unless stated otherwise. Sections were first incubated with a peroxidase blocker (PX968M, BioCare Medical) for 5 minutes. This was followed by three wash steps with wash solution, a 10-minute incubation with background blocker and a further three washes. The HER2 primary antibody (RM-9103-S0, ThermoFisher) was then added to the sections for 15 minutes. The sections were washed three times and then incubated with X-cell HRP-polymer (RHRP520L, BioCare Medical) for 30 minutes. A further three washes were carried out, followed by one wash with deionised water and then a 10-minute incubation with DAB+ substrate chromogen system (K3468, Dako). The sections were washed with deionised water three times, stained with Haematoxylin QS (H-3404, Vector Laboratories) for 1 minute and washed with deionised water another three times. Sections were allowed to dry for 10 minutes before mounting onto coverslips with ClearVue® Mountant XYL mounting medium solution. A secondary control (without primary HER2 antibody) for each sample was also run alongside.

### Immunofluorescence (IF)

The staining protocol for IF was carried out on sections that were mounted onto coverslips instead of glass slides. As detailed above, sections were dewaxed and heat-induced antigen retrieval was carried out. The sections were then washed three times with Leica BOND wash buffer, permeabilised with 0.2% triton X-100 for 15 minutes and then blocked for 1 hour using the Leica BOND background blocker. The blocker was removed and replaced with the primary antibody against the intracellular region of HER2 (1:100, MA1-35720) diluted in Leica BOND antibody diluent for 2 hours. A secondary control (without primary HER2 antibody) for each sample was also run alongside. The primary antibody was washed off the tissue with wash buffer three times followed by 1 hour incubation with the secondary antibody diluted 1:1000 in antibody diluent (goat anti-mouse IgG secondary antibody, Alexa Fluor™ 647, A21236, Invitrogen). The tissue was washed three times with wash buffer, a post-fixation step of 4% paraformaldehyde for 10 minutes was carried out, followed by three more washes before the tissue was stored at 4 °C overnight in phosphate buffered saline (PBS) prior to imaging.

### dSTORM method development

To image tissue sections on coverslips on the Zeiss Elyra7 inverted microscope setup we needed: (1) a chamber that sits on the coverslip that we could add the imaging buffer to, and (2) an adaptor slide, to allow a coverslip to be placed on the current microscope setup. A chamber and adaptor slide were 3D printed using PLA and an Ender 3 Max printer. The chamber was attached to the coverslip using Kwik-Sil™ silicone adhesive or nail polish which set in ∼5 minutes. The chamber was removed from the coverslip after image acquisition using 70% ethanol to soften the adhesive. The coverslips were then mounted onto a microscope slide using Prolong Gold mounting medium (P36934, Invitrogen) for widefield imaging.

### Widefield microscopy

H&E and IHC images were acquired on the Zeiss AxioScan.Z1 with a Plan Apochromat 20x/0.8 M27 objective and Hitachi HV-F203SCL camera. Widefield immunofluorescence of all sections was carried out using the Zeiss Axio Observer 7 and Hamamatsu camera with Plan Apochromat 20x/0.8 M27.

### dSTORM imaging

dSTORM imaging was carried out using a Zeiss Elyra7 setup

A switching buffer was used to provide the optimal environmental conditions for fluorophore switching ^40^. In brief, three solutions were made in advance (A, B and C), Table 4. Immediately prior to imaging, solutions A, B and C were combined with PBS at a ratio of 1:8:2:9, respectively. The sample chamber was filled with switching buffer, and a microscope slide was placed on top to prevent air reaching the buffer.

**Table 4:**
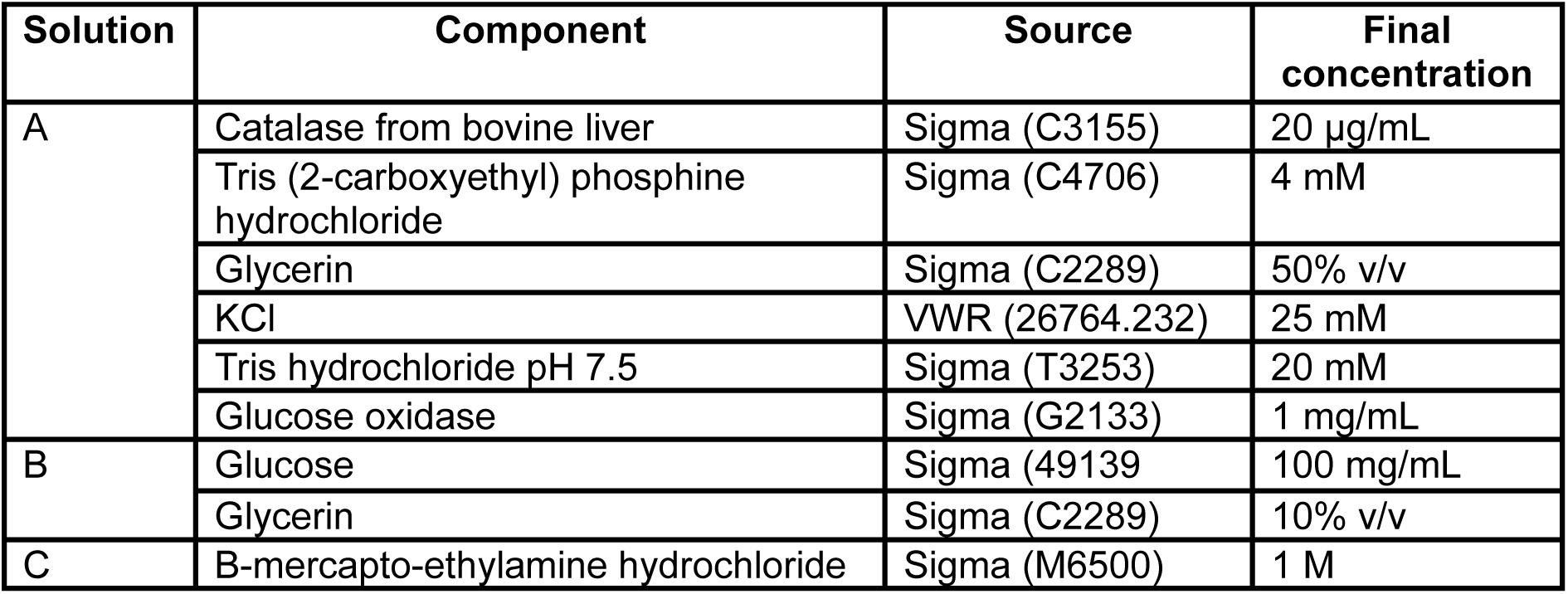
Imaging buffer components for dSTORM imaging.

| Solution | Component | Source | Final concentration |
| --- | --- | --- | --- |
| A | Catalase from bovine liver | Sigma (C3155) | 20 µg/mL |
|  | Tris (2-carboxyethyl) phosphine hydrochloride | Sigma (C4706) | 4 mM |
|  | Glycerin | Sigma (C2289) | 50% v/v |
|  | KCl | VWR (26764.232) | 25 mM |
|  | Tris hydrochloride pH 7.5 | Sigma (T3253) | 20 mM |
|  | Glucose oxidase | Sigma (G2133) | 1 mg/mL |
| B | Glucose | Sigma (49139) | 100 mg/mL |
|  | Glycerin | Sigma (C2289) | 10% v/v |
| C | B-mercapto-ethylamine hydrochloride | Sigma (M6500) | 1 M |

The Zeiss Elyra7 STORM setup was used with alpha Plan-Apochromat 100x/1.46 Oil DIC objective. Samples were exposed to a high laser power briefly to push fluorophores into the dark state. The imaging angle was moved slightly out of total internal reflective fluorescence (TIRF) into a highly inclined laminated optical (HILO) angle. For method development with the PDX samples, 5,000 frames were acquired. For the clinical samples, 10,000 frames were acquired. Images were acquired with an exposure time of 30 ms, high 647 nm laser power and increasing 405 nm laser power to assist with blinking.

Following the acquisition, dSTORM images were reconstructed and filtered using the Zeiss Zen Black software. Filtering was based on the localisation precision (7-35 nm), photon count (>500), background variance, Chi square and point spread function. Localisation tables were exported from the Zen software for analysis. For the PDX samples, 13-20 fields of view were imaged over 2-3 repeats. For the clinical samples, 17-18 fields of view were imaged over 2 repeats.

### Data analysis

The analysis was performed using the principles of literate programming ^41^, combining analysis and documentation, implemented in R ^42^ using quarto^43^. Principles of defensive programming were implemented using the R package checkmate ^44^, the units ^45^ package was used throughout for robust handling of SI units. All tabular data was managed with the package data.table ^46^. The data generated by Zen Black was aggregated into one table for the PDX samples and one for the clinical samples. The packages ggplot2 ^47^, see ^48^ and ggdist ^49^ were used to visualise the clustering output examples, the distributions of the summary measures and the results of the statistical analysis.

### Cluster analysis

The clustering of HER2 localisations was performed using the Topological Mode Analysis Tool (ToMATo) algorithm implemented in the package RSMLM ^29^. The search radius was set to 50 nm and the persistence threshold to 15.

The images illustrating the output of the ToMATo clustering were created by computing the 6 nearest neighbours of each cluster detected (centre to centre). The package igraph ^50^ was then used to ensure good contrast between neighbouring clusters in the resulting images.

The package sf ^51^ was used to estimate the area of the polygon described by localisations in each cluster identified by RSMLM. A concave hull ratio of 0.1 was used to define the polygon area, thus discarding the outer regions of the convex hull with no detected localisations, and a HER2 radius of 7 nm was assumed, in accordance with the hydrodynamic diameter reported by Vicente-Alique *et al.* (2011) ^52^. The output of this process was collated into one table for PDX samples and one for clinical samples. For each dSTORM data set, the number of clusters detected per image was divided by the tissue area in that image determined by Fiji ImageJ ^53,54^ and collated into separate tables.

### Statistical analysis

The number of detected localisations per cluster was analysed using a generalised linear mixed model with the package glmmTMB ^55^, assuming negative binomial distributions. The other summary measures reported were log-transformed and analysed using linear mixed-effects models with the package lmerTest ^56^. In both cases, HER2 grade was set as the fixed effect and STORM image, within tissue section, within biological sample were set as random effects. Tukey’s HSD tests were used for multiple comparisons of means using the package multcomp ^57^. The EMD ^58,59^ was used to compare distributions of summary measures across biological samples, this method was chosen as it is purely data-driven and makes no assumption about the shape of these distributions. The total tissue area, number of localisations and number of clusters used in the analysis for the PDX and clinical samples are summarised in Supplementary Table 1 and Table 2, respectively. Data are reported as the model estimates and 95% confidence intervals.

Supplementary information: Additional results including control data for imaging techniques, HER2 cluster densities, H&E and widefield immunofluorescence of HER2 in clinical samples, comparison of dSTORM images acquired from 5,000 and 10,000 frames and a summary of the data used for analysis (PDF).

## Acknowledgements

We thank P. Auckland (Medicines Discovery Catapult) for helpful discussions and review of the manuscript, G. Ashton (Cancer Research UK, Manchester) for sectioning the clinical tissue and M. Eyres (Medicines Discovery Catapult) for 3D printing of coverslip chamber and holder.

