## Supplementary Figures 1-7, Supplementary Tables 1-2 for "Grading HER2 at the nanoscale in clinical tissue"

### Supporting Information

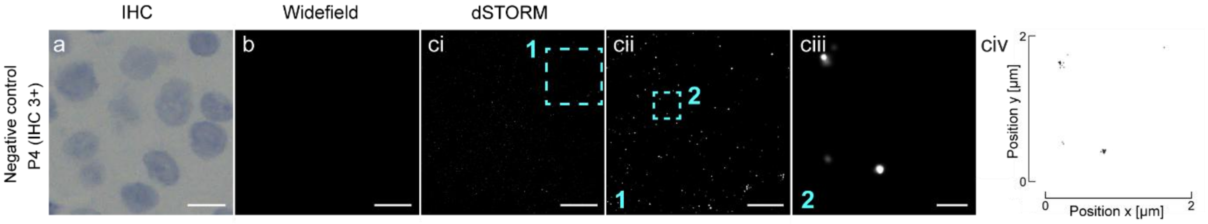


**Supplementary Figure 1: HER2 negative staining control images for each imaging technique.** Representative images for (a) IHC, (b) widefield fluorescence microscopy, and (c) dSTORM. dSTORM images (ci-ciii) are shown with a gaussian render. (cii) and (ciii) show the boxed regions in (ci) and (cii), respectively. Individual localisations in (ciii) are shown in (civ). PDX sample (P4) with IHC HER2 grade of 3+ was used. Scale bars: 10 µm (a, b and ci), 2.5 µm (cii) and 500 nm (ciii).

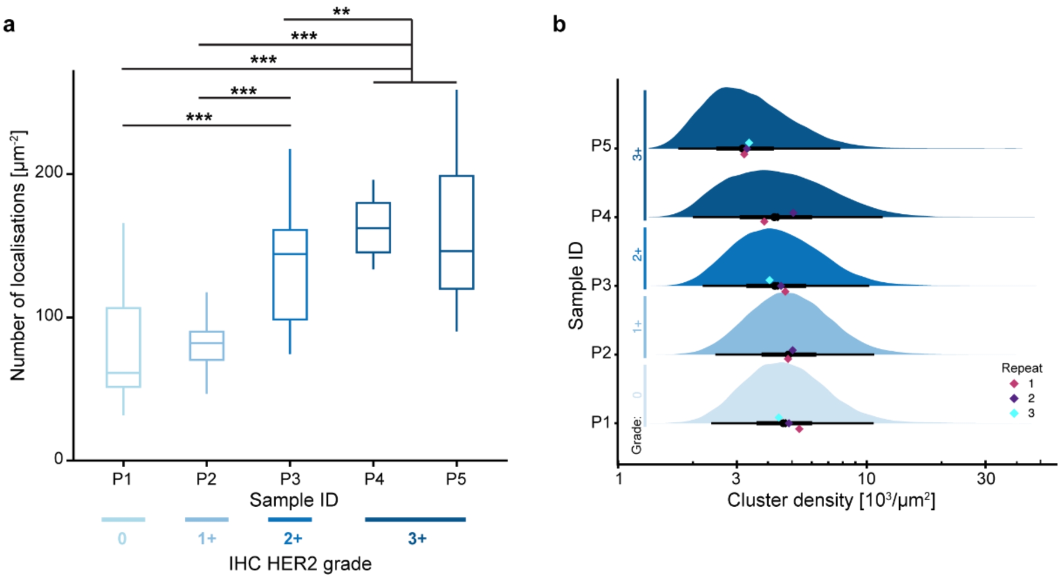


**Supplementary Figure 2: Number of localisations per tissue area (a) and HER2 cluster density (b) in five PDX samples (P1-5).** Cluster density is the number of localisations per cluster area quantified from direct stochastic optical reconstruction microscopy images of HER2. The sample mean (black diamond) and median (black circle) are shown and the lines represent the interquartile range (thick line) and the range of 95% of the data (thin line) (b). N = 2,616,032-8,507,907 localisations and 42,328-87,517 clusters from 3 technical repeats. Significant differences were determined using a Linear Mixed-Effects model with Tukey HSD test. $p \leq 0.01$ (**), $p \leq 0.001$ (***).

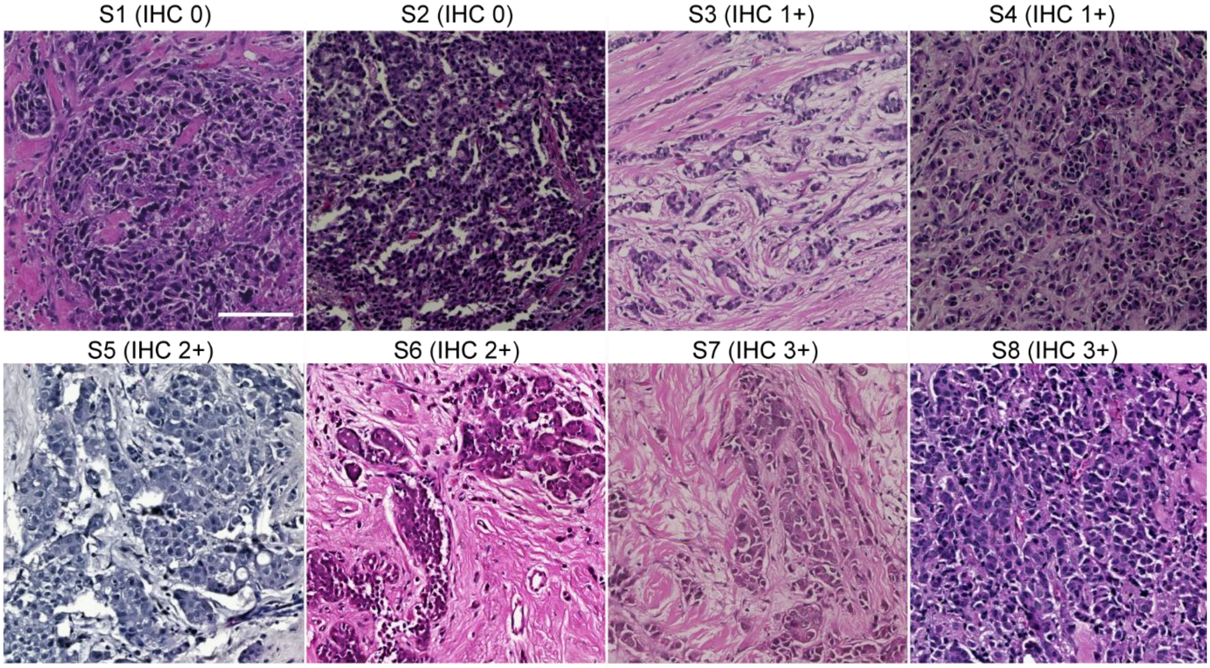


**Supplementary Figure 3: Haematoxylin and Eosin stain of eight clinical breast tumours (S1-S8).** HER2 grading was carried out by a research pathologist using immunohistochemistry. Scale bar:100 μm.

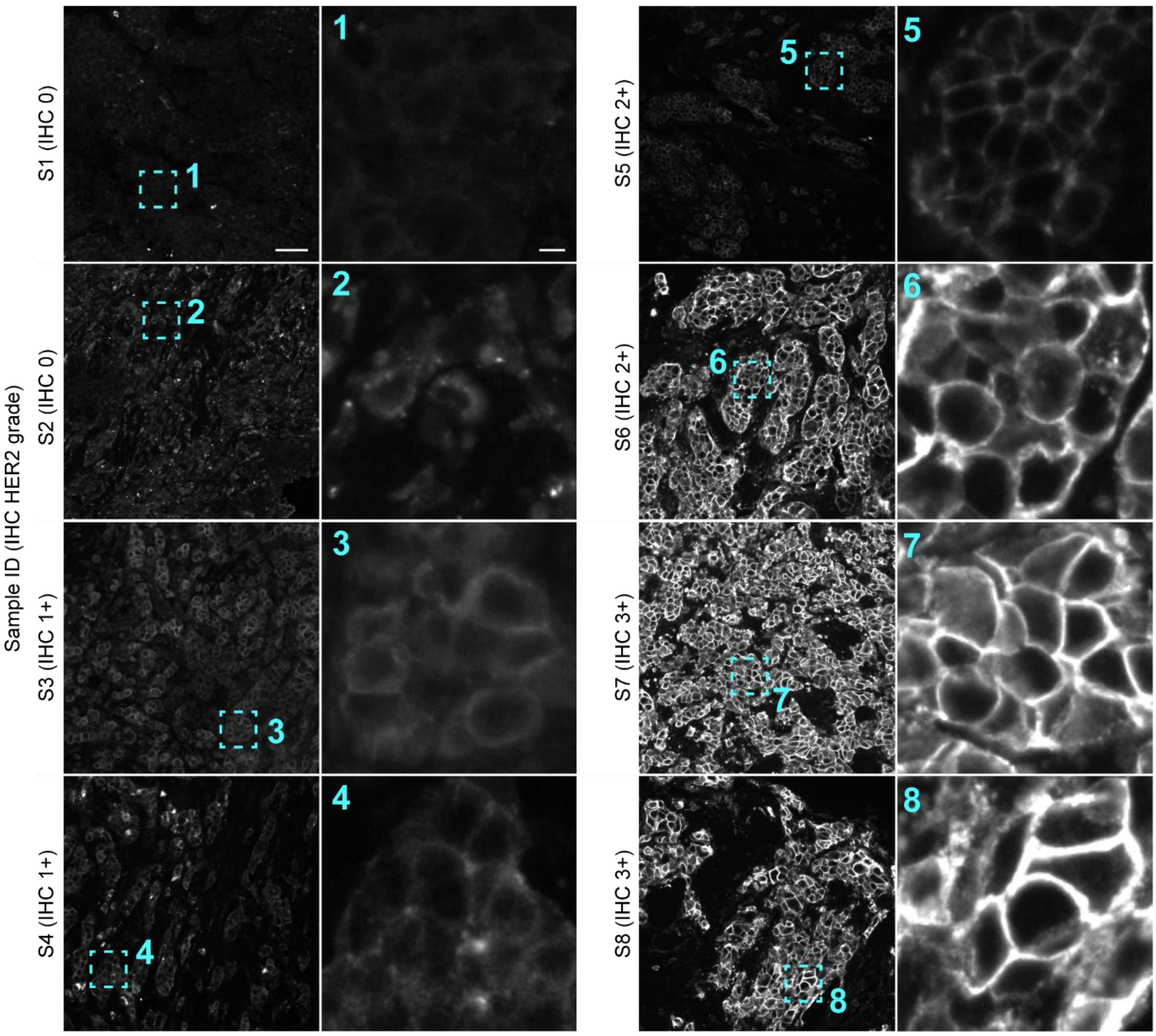


**Supplementary Figure 4: Widefield immunofluorescence images of HER2 labelling of eight clinical breast tumour samples (S1-8).** HER2 grade was determined by a research pathologist using immunohistochemistry (IHC 0-3+). Scale bar: 50 µm and 5 µm (zoomed region).

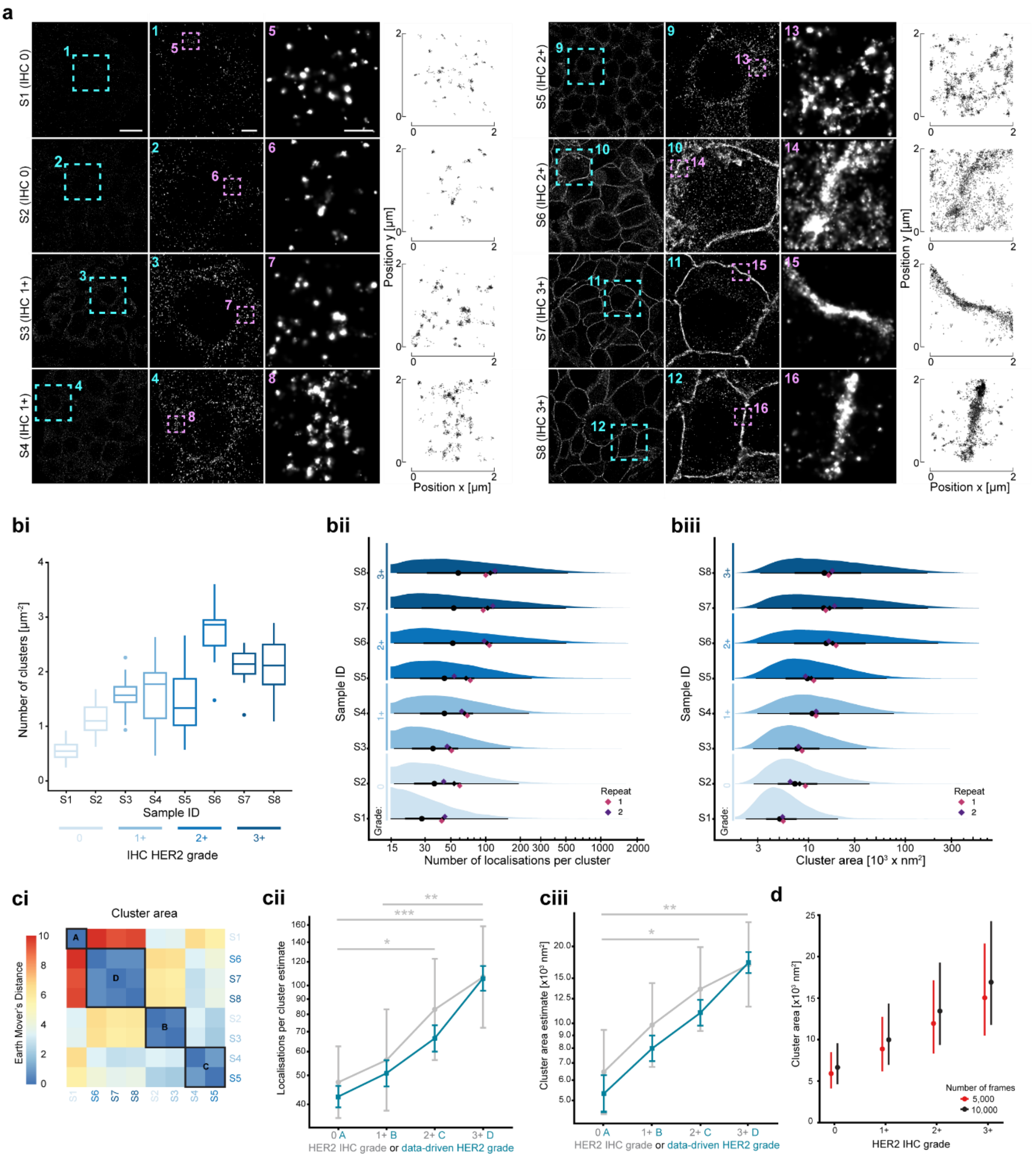

**Supplementary Figure 5: HER2 cluster analysis in eight clinical breast tumour samples (S1-8).** (a) Example direct stochastic optical reconstruction microscopy images of HER2 for each sample using 10,000 frames. Zoomed regions are shown by the boxes and respective numbering. Individual localisations are shown for the second zoomed image. The immunohistochemistry (IHC) grade is stated next to the sample ID. Scale bars (from left to right): 10 µm, 2 µm and 500 nm. The number of HER2 clusters per tissue area (bi), cluster area (bii) and number of localisations per cluster (biii) were quantified. The sample mean (black diamond) and median (black circle) are shown with lines representing the interquartile range (thick line) and the range of 95% of the data (thin line) (bii and biii). N = 22,817-119,756 clusters from 2 technical repeats. The Earth-Mover's Distance heatmap (ci) shows the data-driven grouping of the samples into A, B, C and D. Comparison of the mean ± 95% confidence interval for the IHC grading and data-driven grading is shown in (cii) for number of localisations per cluster and (ciii) for cluster area. (d) Comparison of HER2 cluster area when acquiring 5,000 frames versus 10,000. Significant differences were determined using a Linear Mixed-Effects model with Tukey HSD test. $p \leq 0.05$ (*), $p \leq 0.01$ (**), $p \leq 0.001$ (***). Significant differences between HER2 IHC grades only are shown in cii and ciii.

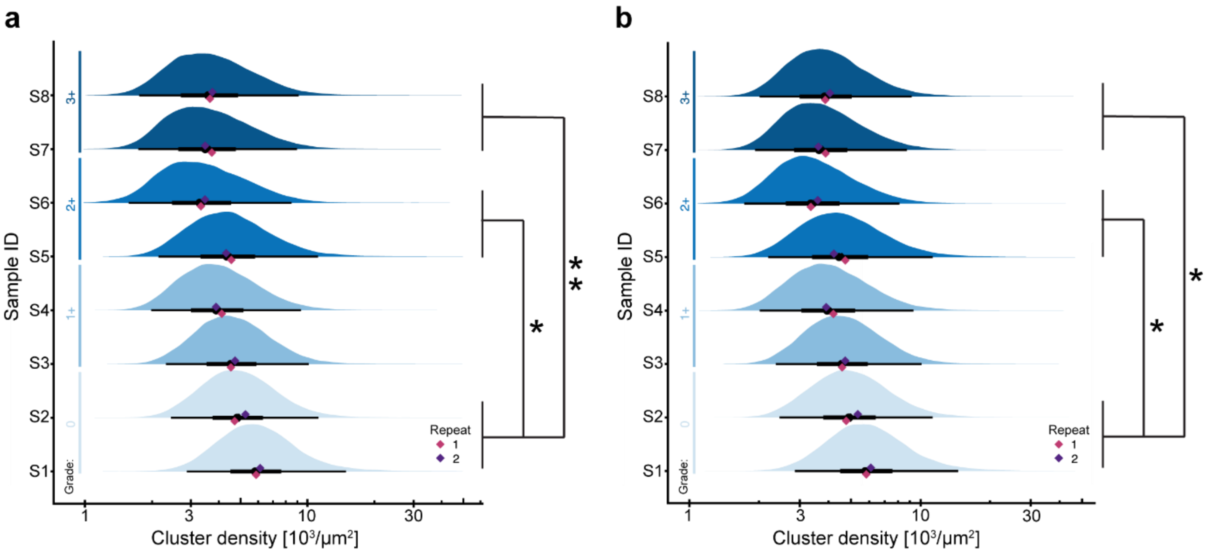


**Supplementary Figure 6: HER2 cluster density in eight clinical breast tumour samples (S1-8).** Cluster density is the number of localisations per cluster area quantified from direct stochastic optical reconstruction microscopy images from 5,000 frames (a) and 10,000 frames (b). The sample mean (black diamond) and median (black circle) are shown with lines representing the interquartile range (thick line) and the range of 95% of the data (thin line). N = 22,817-119,756 clusters from 2 technical repeats. Significant differences were determined using a Linear Mixed-Effects model with Tukey HSD test. $p \leq 0.05$ (*), $p \leq 0.01$ (**).

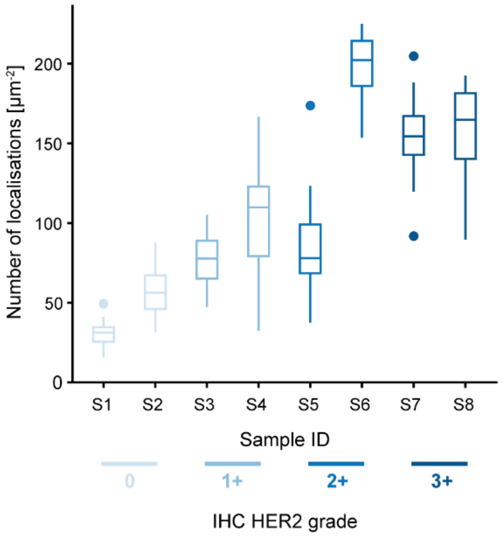


| Comparing grades | P value |
|---|---|
| 0 vs 1+ | < 0.001 |
| 0 vs 2+ | < 0.001 |
| 0 vs 3+ | < 0.001 |
| 1+ vs 2+ | < 0.001 |
| 1+ vs 3+ | < 0.001 |
| 2+ vs 3+ | 0.07 |

**Supplementary Figure 7: Number of detected localisations in eight clinical breast tumour samples (S1-8) from dSTORM images from 5,000 frames.** N = 1,340,090-9,313,711 localisations from 2 technical repeats. Significant differences were determined if $p \leq 0.05$ using an ANOVA with Tukey HSD test.

**Supplementary Table 1: Total tissue area, number of localisations and number of clusters analysed per PDX sample.**

| Sample ID | Tissue area [$\mu m^2$] | Number of localisations | Number of clusters |
|---|---|---|---|
| P1 | 38755.98 | 3080434 | 46254 |
| P2 | 32715.57 | 2616032 | 42328 |
| P3 | 38986.58 | 5335315 | 69462 |
| P4 | 52066.84 | 8507907 | 87517 |
| P5 | 41676.81 | 6735738 | 52415 |

**Supplementary Table 2: Total tissue area, number of localisations and number of clusters analysed per clinical sample for 5,000 and 10,000 frame datasets.**

| Sample ID | Tissue area [$\mu m^2$] | Number of localisations (5,000 frames) | Number of clusters (5,000 frames) | Number of localisations (10,000 frames) | Number of clusters (10,000 frames) |
|---|---|---|---|---|---|
| S1 | 43585.68 | 1340090 | 20197 | 1568918 | 24123 |
| S2 | 45721.08 | 2638470 | 40560 | 3563942 | 52496 |
| S3 | 46544.10 | 3588759 | 54945 | 4777082 | 72702 |
| S4 | 40521.28 | 4139680 | 52896 | 5306051 | 63822 |
| S5 | 45768.29 | 3962296 | 51599 | 5608750 | 67658 |
| S6 | 46885.95 | 9313711 | 83427 | 15870557 | 127392 |
| S7 | 46946.12 | 7236903 | 65311 | 12273392 | 99568 |
| S8 | 44913.55 | 7089683 | 66820 | 11755668 | 95716 |
